# Efficient Gene Knockout in Salivary Gland Epithelial Explant Cultures

**DOI:** 10.1101/2022.02.28.482127

**Authors:** Rei Sekiguchi, Madison M. Mehlferber, Kazue Matsumoto, Shaohe Wang

**Author notes:** Correspondence to: Shaohe Wang. **One-sentence description of our article:** We present a CRISPR-Cas9 mediated gene perturbation system in mouse embryonic salivary epithelial explant cultures and use it to demonstrate a critical role for β1 integrin in branching morphogenesis.

## Abstract

We have developed methods to achieve efficient CRISPR-Cas9 mediated gene knockout in *ex vivo* mouse embryonic salivary epithelial explants. Salivary epithelial explants provide a valuable model for characterizing cell signaling, differentiation, and epithelial morphogenesis, but research has been limited by a paucity of efficient gene perturbation methods. Here, we demonstrate highly efficient gene perturbation by transient transduction of guide RNA-expressing lentiviruses into Cas9-expressing salivary epithelial buds isolated from Cas9 transgenic mice. We first show that salivary epithelial explants can be cultured in low-concentration, non-solidified Matrigel suspensions in 96-well plates, which greatly increases sample throughput compared to conventional cultures embedded in solidified Matrigel. We further show that salivary epithelial explants can grow and branch with FGF7 alone, while supplementing with ITS (insulin, transferrin, and selenium) enhances growth and branching. We then describe an efficient workflow to produce experiment-ready, high-titer lentiviruses within 1 week after molecular cloning. To track transduced cells, we designed the lentiviral vector to co-express a nuclear fluorescent reporter with the guide RNA. We routinely achieve 80% transduction efficiency when antibiotic selection is used. Importantly, we detected robust loss of targeted protein products when testing 9 guide RNAs for 3 different genes. Moreover, targeting the β1 integrin gene (*Itgb1*) inhibited branching morphogenesis, which supports the importance of cell-matrix adhesion in driving branching morphogenesis. In summary, we have established a lentivirus-based method that can efficiently perturb genes of interest in salivary epithelial explants, which will greatly facilitate studies of specific gene functions using this system.

## INTRODUCTION

The mouse salivary gland is an excellent model for studying mammalian organ development. During mouse embryogenesis, an epithelial bud bulges from the oral epidermis to become surrounded by neural crest-derived mesenchyme to form the primordial salivary gland (Jaskoll et al. 2002). Between the epithelium and mesenchyme is a dense, thin layer of extracellular matrix, the basement membrane (Sekiguchi and Yamada 2018). The initial epithelial bud grows and repetitively splits into numerous buds arranged in a “bunch of grapes” appearance through branching morphogenesis, while epithelial cells differentiate into ductal, acinar, or myoepithelial cells, whose concerted actions enable saliva secretion in matured organs (Patel et al. 2006; Tucker 2007; Wang et al. 2017).

Salivary gland research has been advanced by a combination of *in vivo* and *ex vivo* approaches. Mouse gene knockout studies definitively established the importance of various signaling pathways, transcriptional factors, and extracellular matrix regulation in salivary gland development (Patel et al. 2006; Knosp et al. 2015; Chatzeli et al. 2017; Emmerson et al. 2017; Szymaniak et al. 2017; Athwal et al. 2019). On the other hand, the *ex vivo* organotypic culture of mouse embryonic salivary glands enabled characterizations and manipulations that are only possible outside embryos (Borghese 1950). For example, live imaging of *ex vivo* cultured salivary glands revealed extensive dynamics of epithelial cells and the basement membrane matrix, which catalyzed the discovery that a combination of strong cell-matrix and weak cell-cell adhesions in surface epithelial cells drives branching morphogenesis of salivary glands (Larsen et al. 2006; Wei et al. 2007; Harunaga et al. 2014; Wang et al. 2021). Furthermore, the mesenchyme encapsulating the salivary epithelium can be surgically removed to allow mesenchyme-free culture of the salivary epithelium in the presence of basement membrane extract and certain growth factors (Nogawa and Takahashi 1991). These salivary epithelial explants enabled analysis of how different growth factors affect their growth, morphogenesis, or differentiation (Steinberg et al. 2005; Nakao et al. 2017).

Small-molecule inhibitors, blocking antibodies, soluble dominant-negative receptors, and small interfering RNA (siRNA) have all been used to perturb gene functions in *ex vivo* cultures of salivary glands (Sakai et al. 2003; Steinberg et al. 2005; Wei et al. 2007), although each has some limitations. Small-molecule inhibitors can be effective to perturb certain signaling pathways or enzymatic activities, but their performance varies depending on their efficacy and specificity. Blocking antibodies or soluble dominant-negative receptors can selectively inhibit ligand binding to surface receptors, but they are only available for a small number of targets. While siRNA-mediated gene knockdown can generalize to most genes, it comes with many disadvantages including transient inhibition, unpredictable knockdown levels, and significant off-target effects.

Compared to siRNA-mediated gene knockdown, CRISPR-Cas9 mediated gene knockout offers persistent disruption and better specificity (Boettcher and McManus 2015). CRISPR-Cas9 is an RNA-guided DNA endonuclease that can make site-specific double-stranded DNA breaks in mammalian cells (Jinek et al. 2012; Cong et al. 2013). These DNA breaks trigger endogenous DNA repair machinery to catalyze non-homologous end joining (NHEJ) or homology-directed repair (HDR), which can be leveraged for various purposes of genome editing. Notably, knockout of protein coding genes can be readily achieved by frameshift mutations introduced by error-prone NHEJ repair. CRISPR-Cas9 has not been used in *ex vivo* salivary gland cultures, partly due to a lack of an efficient delivery system for the single guide RNA (sgRNA) needed to target a specific gene.

Here, we demonstrate highly efficient CRISPR-Cas9 mediated gene knockout in *ex vivo* cultures of mouse embryonic salivary epithelial explants, which was achieved by transient transduction of sgRNA-expressing high-titer lentiviruses into Cas9-expressing salivary epithelial buds isolated from Cas9 transgenic mice (Platt et al. 2014). To facilitate application of this method, we developed new isolation and culture techniques for salivary epithelial explants that greatly improved sample throughput, and an efficient workflow to produce experiment-ready, high-titer lentiviruses within 1 week after molecular cloning. Importantly, we showed that branching morphogenesis was inhibited when the β1 integrin gene (Itgb1) was targeted, indicating our lentiviral delivery system is efficient enough to produce loss-of-function phenotypes at the tissue level. This approach has broad applications to test the roles of genes that may contribute to salivary gland development or disease.

## MATERIALS AND METHODS

### Mouse strains

Mouse experiments were approved by the NIDCR Animal Care and Use Committee (Animal Study Protocols 17-845 and 20-1040). Standard mouse housing and husbandry were provided by the NIDCR Veterinary Resource Core. Timed pregnant ICR (CD-1) outbred mice were obtained from Envigo to get wildtype embryos. The Cas9 transgenic mice (Platt et al. 2014) were obtained from the Jackson Laboratory (JAX, 026558). To generate embryos at specific gestational stages, transgenic mice 8-16 weeks old were bred, where the next day after a vaginal plug was found was defined as embryonic day 1.

### Experiment design

This study complied with Animal Research: Reporting In Vivo Experiments (ARRIVE) 2.0. For each experiment, all submandibular salivary glands from all embryos without sex identification (mixed sex) from one timed pregnant mouse were used to isolate individual epithelial buds. The epithelial buds with similar sizes were selected for transduction with lentiviruses expressing a control sgRNA or an sgRNA targeting a gene of interest. The extent of branching morphogenesis or the targeted protein expression level was compared between explants grown from epithelial buds expressing the control sgRNA and those expressing the targeting sgRNA. No animals or embryos were excluded from experiments. Criteria used for excluding qPCR data points were detailed in the qPCR section. Exact sample sizes are provided in the figure legends.

### sgRNA design

The first half of the coding sequence of a target gene was used for sgRNA design using CRISPOR (http://crispor.tefor.net/) (Concordet and Haeussler 2018) with the UCSC mm10 mouse reference genome and the NGG protospacer adjacent motif. Three sgRNAs with both high specificity and efficiency scores were selected for each target gene (Hsu et al. 2013; Bae et al. 2014; Doench et al. 2016; Chen et al. 2019). sgRNAs with motifs predicted to lower gene knockout efficiencies were avoided (Graf et al. 2019). An extra G was added to the 5’-end if the sequence did not begin with G to facilitate transcription by the U6 promoter.

### Plasmids

The parental lentiviral vector (pW212; Addgene, 170810) co-expressing sgRNA with an red nuclear fluorescent reporter (NLS-mScarlet-I) and a blasticidin resistance gene was previously generated by our group (Wang et al. 2021). To make the parental vector (pW299; **Addgene, TBD**) with a blue nuclear fluorescent reporter (NLS-tagBFP), mScarlet-I in pW212 was replaced with tagBFP using Gibson Assembly (Gibson et al. 2009). Two synonymous mutations were introduced to tagBFP to remove an Esp3I site. For cloning of lenti-sgRNA plasmids, a pair of oligos with the desired sgRNA sequence (see **sgRNA design**) plus a 4-bp 5’-extension (“cacc” for the forward oligo and “aaac” for the reverse complementary oligo) were annealed, and the resulting oligo duplex and lentiviral vectors (pW212 or pW299) digested by Esp3I (NEB, R0734S) were ligated using a 1:2 mixture of T4 ligase (NEB, M0202L) and T4 polynucleotide kinase (NEB, M0236L) in T4 ligase buffer. The ligation mix was transformed using NEB stable competent cells (NEB, C3040). Correct insertion of sgRNA sequence was confirmed by Sanger sequencing.

### Lentivirus packaging

Lentivirus packaging was performed as previously described (Wang et al. 2021). Briefly, lenti-sgRNA plasmids were co-transfected with psPAX2 (Addgene, 12260) and pMD2.G (Addgene, 12259) into HEK293T cells by calcium co-precipitation to produce infectious lentiviral particles. Pooled lentivirus-containing media collected at 36- and 60-hours post transfection were passed through a 0.45 µm filter (MilliporeSigma, SE1M003M00), and concentrated using PEG (System Biosciences, LV825A-1) following the manufacturer’s instructions. Virus titer was estimated using Lenti-X GoStix Plus (Takara, 631281) after 100x dilution, where the band intensities were quantified using a smartphone app following the manufacturer’s instructions. Concentrated lentiviruses were stored at -80°C.

### Lentivirus transduction of salivary epithelial buds

Lentivirus transduction was performed in the wells of an ultra-low attachment 96-well V-bottom plate (S-bio, MS-9096VZ). Each epithelial bud was transferred into one well in precisely 5 µL medium. A 15 µL lentivirus treatment mixture containing 10 µL lentivirus stock, 4 µL DMEM/F- 12 and 1 µL 160 µg/mL polybrene (MilliporeSigma, H9268) was added to each well. The plate was incubated in a humidified 37°C incubator for 1-2 hours. Each bud was washed 3 times in DMEM/F-12 before culture (see **Explant culture**).

### Salivary epithelial bud isolation

Mouse submandibular salivary glands were isolated from 13 or 13.5 day embryos as previously described (Sequeira et al. 2013). The attached sublingual gland was removed by dissection after each gland was isolated to ensure all epithelial buds were from submandibular glands. Glands were treated with 2 units/mL dispase (Thermo Fisher, 17105041; diluted in DMEM/F-12) in a Pyrex spot plate (Fisher Scientific 13-748B; 6-10 glands per well) for 15 min at 37°C. To quench dispase activity, glands were washed twice with 5% BSA (w/v; MilliporeSigma, A8577; diluted in DMEM/F-12). While being monitored under a dissecting microscope, glands were repetitively triturated using a 200 µL pipettor set at 100 µL with a low-retention tip (Rainin, 30389187), until the mesenchyme was dissociated into single cells while epithelial buds remained intact. Salivary epithelial buds were rinsed 3 times by being transferred to new wells of the spot plate prefilled with 150 µL 5% BSA (w/v) in DMEM/F-12 using a 20 µL pipettor with a low-retention pipette tip (Rainin, 30389190). Care was taken during transfer to minimize carryover of mesenchymal cells. To prevent evaporation, the well was covered with a glass coverslip during preparation of lentivirus transduction (see **Lentivirus transduction of salivary epithelial buds**) and explant culture media (see **Explant culture**). Before the next steps, epithelial buds were further rinsed twice in DMEM/F-12 without BSA.

### Explant culture

Before salivary gland dissections, aliquots of growth factor-reduced Matrigel (Corning, 356231; stock 9-10 mg/mL) were thawed at 4°C or on ice. The volume was calculated as 5 µL per explant with 1 or 2 µL extra to compensate for pipetting errors. The base medium for culture was DMEM/F-12 (Thermo Fisher, 11039047) supplemented with 1x PenStrep (100 units/mL penicillin, 100 µg/mL streptomycin; Thermo Fisher, 15140163), hereafter referred to as DMEM/F- 12-PS.

Epithelial buds were cultured in ultra-low attachment 96-well V-bottom plates (S-bio, MS- 9096VZ) as previously described (Wang et al. 2021). Briefly, one bud was cultured in each well in the explant culture media with 0.5 mg/mL growth factor-reduced Matrigel (Corning, 356231; stock 9-10 mg/mL), 200 ng/mL FGF7 (R&D Systems, 5028-KG-025), and 1x ITS supplement (10 mg/L insulin, 5.5 mg/L transferrin, 6.7 µg/L selenium; Thermo Fisher, 41400045) in DMEM/F-12- PS. 10 ng/mL NRG1 (R&D Systems, 9875-NR-050) was included in some testing culture conditions. In practice, the explant culture media was prepared at 2x concentration with a total volume of (n + 2) x 50 µL, where n is the sample number. The wells for explant culture were pre-filled with 47 µL DMEM/F-12-PS, and one bud was transferred into each well in precisely 3 µL medium using a low-retention pipette tip. 50 µL 2x explant culture media was then added to each well. The plate was cultured at 37°C with 5% CO_2_.

### Quantitative PCR (qPCR) of explants

Total RNA from each explant culture was extracted using the RNeasy Micro Kit (Qiagen, 74004) following the manufacturer’s instructions and also described here. 10 µL β-ME (MilliporeSigma, M3148) was added per 1 mL Buffer RLT (lysis buffer). Each explant was dissolved in 100 µL Buffer RLT by vortexing and stored at -80°C until further processing. Once the sample was thawed, 100 µL of 70% ethanol was added and mixed by pipetting. The mixture was transferred onto the spin column and centrifuged for 15 s at 8,000x g. To wash the spin column, 350 µL Buffer RW1 was added and centrifuged for 15 s at 8,000x g. To digest genomic DNA, 80 µL DNase I mix (10 µL DNase I stock + 70 µL Buffer RDD) was added to the column and incubated for 15 min at room temperature. 350 µL Buffer RW1 was added and centrifuged for 15 s at 8,000x g to wash the spin column. To further wash the spin column, it was placed in a new collection tube and washed sequentially with 500 µL Buffer RPE (centrifugation for 15 s at 8,000x g) and 500 µL 80% ethanol (centrifugation for 2 min at 8,000x g). To dry the spin column, it was placed in a new collection tube and centrifuged at full speed for 5 min. To elute RNA, 14 µL RNase-free water was added directly to the center of the spin column membrane and centrifuged at full speed for 1 min with the lid closed.

To synthesize cDNA from total RNA, we used the iScript cDNA Synthesis Kit (Bio-Rad, 1708891). A 20 µL reaction containing 10 µL total RNA, 5 µL nuclease-free water, 4 µL 5x iScript reaction mix, and 1 µL iScript reserve transcriptase was incubated in a thermo cycler with the following: 5 min at 25°C (priming), 20 min at 46°C (reverse transcription), and 1 min at 95°C (reserve transcriptase inactivation). The cDNA reactions were diluted 1:2 with nuclease-free water and stored at -20°C. The diluted cDNA reaction was used as qPCR templates.

qPCR primers were designed using the online primer design tool, Primer 3 (Koressaar and Remm 2007; Untergasser et al. 2012; Kõressaar et al. 2018). To design qPCR primers that can distinguish CRISPR/Cas9-induced mutations from wild type mRNAs, one primer was deigned to overlap the sgRNA cut site, so that there are 3 or 4 bases on the primer’s 3’-end extending beyond the cut site (Li et al. 2019). All qPCR primers used in this study can found in **Appendix Table 2**.

qPCR was performed using the Bio-Rad CFX96 Real-Time System. Each 10 µL qPCR reaction contains 3.5 µL nuclease-free water, 1 µL template (1:2 diluted cDNA reaction), 0.5 µL primer mix (10 µM each in stock, 500 nM each in final reaction), and 5 µL 2x SsoAdvanced Universal SYBR Green Supermix (Bio-Rad, 1725272). 3 technical replicates were performed for each sample/primer combination. For each qPCR assay, a master mix containing primers but not the template was made and aliquoted to 96-well PCR plates (Bio-Rad, HSP9601). Right before loading, the cDNA templates were added to the plate, which was sealed with a sealing film (USA Scientific, 2921-7810), vortexed to mix well, and spun at 100x g for 3 min in a swing-bucket centrifuge (e.g., Eppendorf 5804R). The cycling conditions are 95°C for 3 min, 40x (95°C for 10 sec, 60°C for 20 sec, plate read), followed by melting curve analysis with 0.5°C per cycle from 65 to 95°C (0.5°C/sec ramp).

### qPCR data analysis

The qPCR data was analyzed using the Bio-Rad CFX Maestro Software and customized Python scripts (see **Data, Code and Resource Availability**). The quantification cycle (Cq) was called within the regime of exponential amplification using the Bio-Rad CFX Maestro Software. The Cq values, amplification data, and melting curve data were exported to csv files from the Bio-Rad CFX Maestro Software. The exported csv files were annotated, computed, and combined using customized Python scripts for plotting. For each sample, the difference of the Cq between testing primers (Cq_test) and reference primers (Rps29; Cq_ref) was used to calculate the expression level relative to the reference mRNA: rel_exp = 2^-(Cq_test-Cq_ref)^. The relative expression values were normalized by dividing the average value of the control group before plotting.

Several quality-control criteria were used to exclude certain data points. First, data points with melting curves with more than one peak or a very broad peak were excluded due to putative non-specific amplification. Second, data points with a reference Cq value > 25 were excluded due to unusually low levels of input cDNA (typical reference Cq ranges 19 to 22). Third, outliers of technical replicates were removed, where outliers were defined as the value that makes the relative standard deviation (standard deviation divided by the mean) of the group larger than 25%.

### Immunostaining of explants

Immunostaining of explants was performed as previously described (Wang et al. 2021). Briefly, explants were fixed in 4% PFA (w/v; Electron Microscopy Sciences, 15710) in PBS for 1 hour at room temperature (RT) or overnight at 4°C, permeabilized in PBSTx (PBS with 0.2% v/v Triton-X-100; Thermo Fisher, 28314) for 30 min at RT, blocked in 5% donkey serum (v/v; Jackson ImmunoResearch, 017-000-121) in PBSTx for 2 hours at RT, incubated in primary antibodies diluted in 5% donkey serum (v/v) in PBSTx for 2 days at 4°C, washed 4x 15 min each in PBSTx at RT, incubated in secondary antibodies diluted in PBSTx for 2 days at 4°C, washed 4x 15 min in PBSTx at RT, rinsed once in PBS, and mounted in antifade mountant (Thermo Fisher, P36930) supported by two layers of imaging spacers (Grace Bio-labs, 654004). Immunostaining of β1 integrin used Atto-647N-labeled primary antibodies with 7-day or 8-day incubation at 4°C without secondary antibody staining. All incubation was performed in sample baskets (Intavis, 12.440) in a 24-well plate.

The following antibodies and concentrations were used for immunostaining. Primary antibodies: anti-HSP47 (MilliporeSigma, HPA029198), 1 µg/mL; anti-α3 integrin (R&D Systems, AF2787), 1 µg/mL; anti-α6 integrin (BD Biosciences, 555734), 2 µg/mL; anti-α9 integrin (R&D Systems, AF3827), 1 µg/mL; Atto-565-labeled Hamster anti-β1 integrin (clone Ha2/5, BD Biosciences 555002), 10 µg/mL; anti-collagen IV (MilliporeSigma, AB769), 2 µg/mL. Secondary antibodies: Alexa Fluor 647-labeled donkey anti-rat (Jackson ImmunoResearch, 712-606-150), donkey anti-rabbit (Jackson ImmunoResearch, 711-606-152) or donkey anti-goat (Jackson ImmunoResearch, 705-606-147) were used at 1:200 (1.5-3 µg/mL).

### Microscopy

Phase contrast and epifluorescence images were acquired using a Nikon 10x, 0.3 NA, Plan Fluor objective on a Nikon Ti-E brightfield microscope system with a Hamamatsu Orca Flash 4.0 V3 sCMOS camera controlled by Nikon NIS-Elements software. The JOBS module of the software was used to automatically go through multiple wells in a 96-well plate.

Immunofluorescence images were acquired by laser scanning confocal microscopy using Nikon 20x, 0.75 NA or 40x, 1.3 NA Plan Fluor objectives on a Nikon A1R Confocal Microscope System controlled by Nikon NIS-Elements software, or a Zeiss 63x, 1.4 NA Plan Apo objective on a Zeiss LSM 880 system controlled by Zeiss ZEN software.

### Image processing, analysis and quantification

Image processing, analysis and quantification were mostly performed using Fiji (Schindelin et al. 2012). Cell segmentation was performed using Cellpose (Stringer et al. 2021). Customized ImageJ Macro and Python scripts were used for automating or facilitating image analysis and data visualization (see **Data, Code and Resource Availability**). Before plotting of fluorescence intensity, all raw intensity values were normalized to the average intensity of the control group.

To facilitate simultaneous display of both bright and dim nuclei, the NLS-mScarlet images in all figures were subjected to a gamma of 0.5. The immunofluorescence images for comparison were all acquired and scaled with identical parameters.

The HSP47 intensity was quantified as the mean intensity in the epithelial bud subtracting the background intensity inside the nuclei of the interior bud. The epithelial bud region of interest (ROI) was segmented using the EGFP channel, which was smoothened with a Gaussian filter (r = 5 pixels) and binarized using the “Huang” thresholding algorithm. The nuclear ROI was segmented using the DAPI channel, which was smoothened with a Gaussian filter (r = 1 pixels), background subtracted with a rolling ball method (r = 50 pixels), and binarized using the default thresholding algorithm. The interior bud ROI was obtained by intersecting the nuclear ROI with the epithelial bud ROI shrunken by 20 microns. For each imaging stack, the average measurement of all z slices was calculated as the final HSP47 intensity.

The α3, α6, α9, β1 integrin, and collagen IV intensities were quantified as the peak intensity along a 3 µm width line across the edge of two transduced cells (α3, α9 and β1 integrin) or across the basement membrane next to a transduced cell (α6 integrin and collagen IV), subtracting background intensity that is the average intensity of the beginning 10% of the line length. Five edges were measured on each image from 3-7 explants per experimental group.

Manual counting of bud number was performed on phase contrast images. File names were scrambled by S.W. before counting for observer blinding. The counting by two observers (R.S. and K.Z.) were averaged to minimize personal bias. S.W. decoded and plotted the counting results.

All statistical comparisons were performed using Tukey’s HSD (honestly significant difference) test (for comparisons involving more than 2 groups) or Student’s t-test (for comparisons between 2 groups) with the Python packages statsmodels and scipy.

### Data, Code and Resource Availability

All data of this study are available in Figshare (link to update upon publication). All plasmids are available in Addgene (link to update upon publication). Customized scripts and usage instructions are available from GitHub (link to update upon publication). Selected step-by-step protocols can be found at https://snownontrace.github.io/.

## RESULTS

### Salivary epithelial bud isolation and explant culture in 96-well plates

We first optimized the isolation and culture techniques for salivary epithelial explants to increase sample throughput (**Appendix Fig. 1A**). Conventionally, salivary gland mesenchyme is removed from the epithelium by dissection after dispase treatment. The whole epithelial rudiment or individual epithelial buds are then embedded in high-concentration Matrigel (a basement membrane extract) that quickly solidified. In our new procedure, multiple salivary glands were triturated in bulk after dispase treatment to dissociate the mesenchyme into single cells, whereas the epithelial buds remained intact, presumably due to stronger cell-cell adhesion. Each epithelial bud was then cultured in non-solidified Matrigel in the well of an ultra-low-attachment 96-well plate, which greatly facilitated sample handling and high-throughput imaging using microscopy systems with well-scanning capabilities.

Next, we tested several combinations of growth factors using the new culture format. We found that epithelial explants grew well with FGF7, NRG1 and ITS (insulin, transferrin, and selenium; **Fig. 1A-B**). The explants failed to grow with only NRG1 and ITS, supporting the critical role of FGF signaling in salivary epithelial development (Steinberg et al. 2005). Removing ITS from the combination significantly slowed explant growth, whereas removing NRG1 did not affect explant growth or budding morphogenesis (**Fig. 1A-B**). Thus, we chose to use the FGF7 and ITS combination for standard explant cultures.

**Figure 1.**
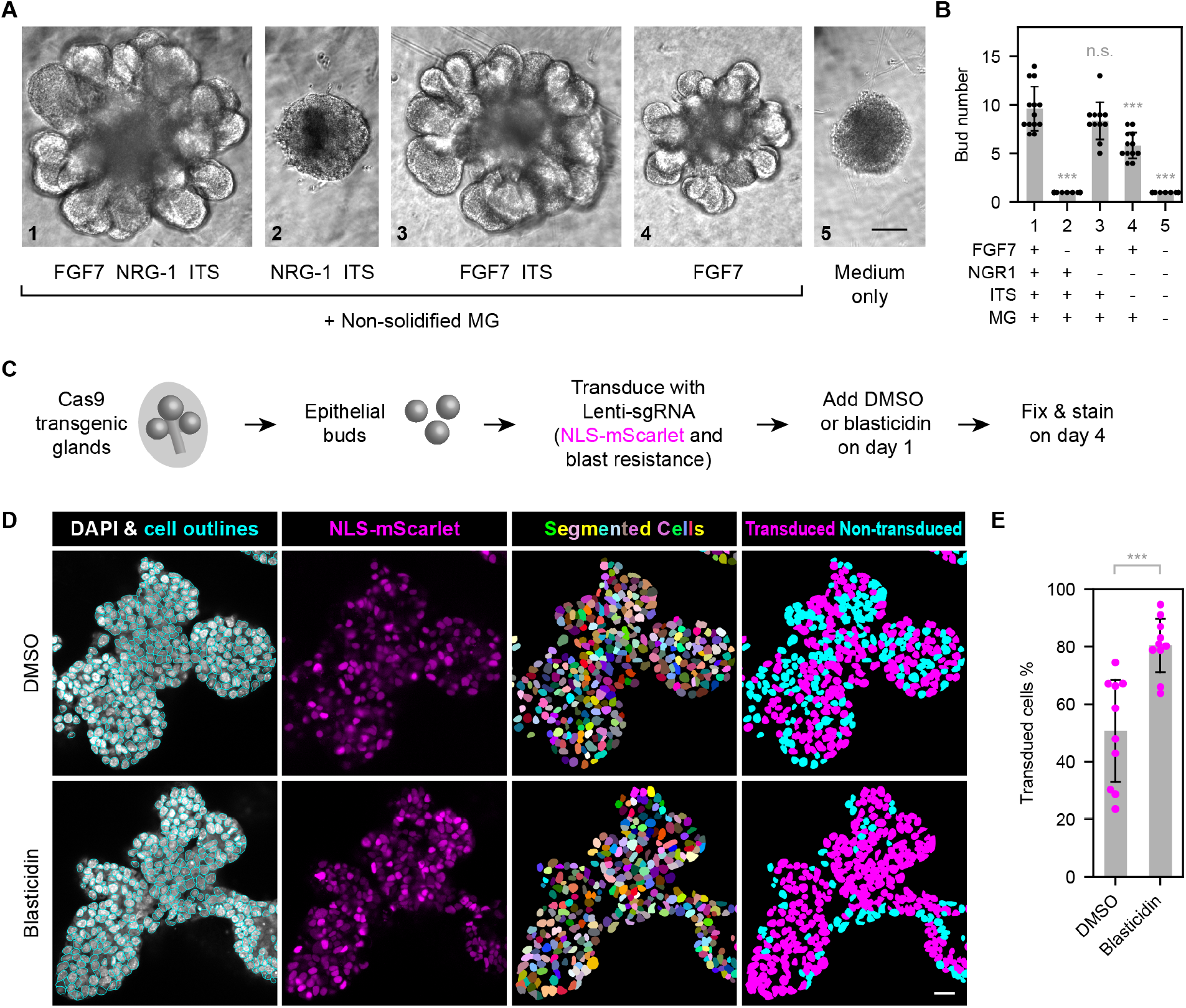
Salivary epithelial explant culture and antibiotic enrichment of lentivirus- transduced cells. (**A**) Phase contrast images of single-bud explant cultures at 2 days under indicated culture conditions. FGF7, 200 ng/mL. NRG1, 10 ng/mL. ITS, 10 mg/L insulin, 5.5 mg/L transferrin, and 6.7 µg/L selenium. Non-solidified Matrigel (MG), 400-500 µg/mL. (**B**) Plot of bud number per explant under indicated culture conditions. Sample number: n = 13, 15, 11, 12, 10 for conditions 1-5. (**C**) Schematics of the lentivirus transduction experiment. (**D**) Nuclear staining and segmentation of explant cultures treated with DMSO (vehicle control) or 20 µg/mL blasticidin. The NLS-mScarlet channel was subjected to a gamma of 0.5 to facilitate simultaneous display of both bright and dim nuclei. (**E**) Plot of percentage of transduced cells in DMSO or blasticidin treated explant cultures at 4 days. Error bars, standard deviation. Statistics, Tukey test (B) or Student’s t-test (E). ***, p<0.001. n.s., not significant. Scale bars, (A) 100 µm, (D) 20 µm.

### Streamlined production of high-titer sgRNA-expressing lentiviruses

To maximize efficiency, we tested the use of lentivirus to express sgRNAs in salivary epithelial explants. We used a previously published lentiviral vector system (Wang et al. 2021) to co-express a nuclear fluorescent reporter with the sgRNA (NLS-mScarlet-I or NLS-tagBFP) to follow transduced cells, and a blasticidin resistance gene (BlastR) to allow antibiotic selection (**Appendix Fig. 1B**). These vectors also enabled straightforward molecular cloning by one-step ligation of the sgRNA oligo duplex (**Appendix Fig. 1B**), which could be easily adapted for parallel cloning of tens of sgRNA vectors. Using miniprep-grade plasmid DNAs, we generated experiment-ready, high-titer lentiviruses in 5 days (**Appendix Fig. 1C**). To estimate lentivirus titer, we used a quick commercial test to measure the test-band intensity (**Appendix Fig. 1D**) and compared that to a reference virus. The typical titer of sgRNA-expressing lentiviruses using this method was about 1.5×10^8^ IFU/mL. After transient transduction of lentiviruses into epithelial buds, nuclear reporter expression could be detected by the next day and reached a maximum level within two days. About 50% of the cells in the explants expressed nuclear reporters without antibiotic selection, while this ratio is raised to 80% with antibiotic selection (**Fig. 1C-E**), indicating that antibiotic selection can efficiently enrich transduced cells.

### Knockout of β1 integrin inhibits branching morphogenesis

For gene perturbation, we transduced sgRNA-expressing lentiviruses into Cas9- expressing salivary epithelial buds isolated from Cas9 transgenic E13 mouse embryos (Platt et al. 2014). We used an sgRNA targeting the *E. coli* lacZ gene as the control and tested 3 targeting sgRNAs for *Itgb1* (**Appendix Table 1**), which encodes β1 integrin that mediates cell-matrix adhesion (Kechagia et al. 2019). All sgRNA sequences were verified by Sanger sequencing (**Appendix Fig. 2A**).

To use quantitative PCR (qPCR) to distinguish wild type and mutant *Itgb1* mRNAs, we designed qPCR primer pairs with one primer overlapping the CRISPR/Cas9 cut site using an established method (**Fig. 2A**; Li et al. 2019). These primers can efficiently and specifically amplify their targets (**Appendix Fig. 2B**). We found that all 3 *Itgb1* sgRNAs could efficiently induce mutations of the targeted mRNAs, as shown by the reduced levels of wild type mRNAs (**Fig. 2B**).

**Figure 2.**
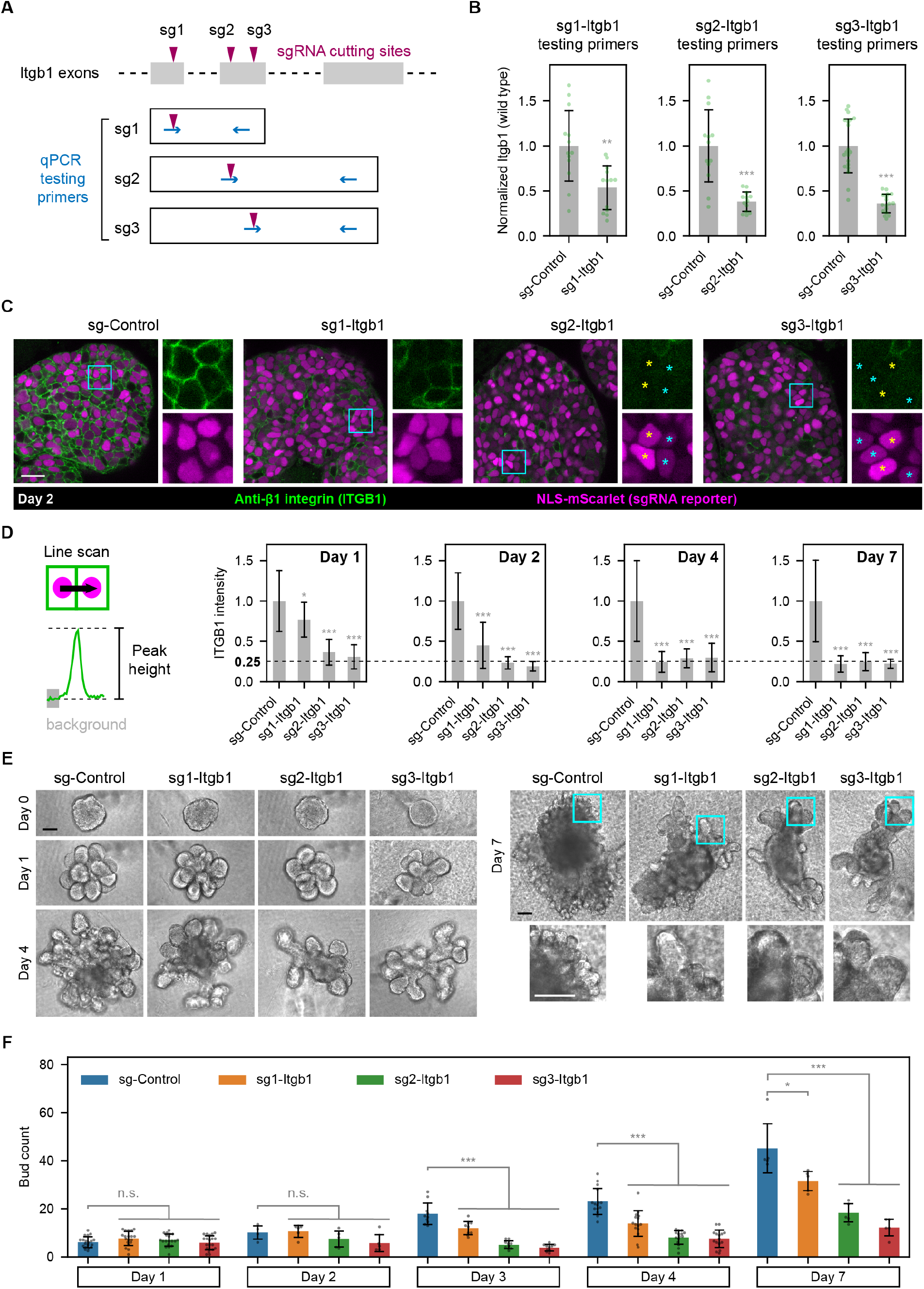
Knockout of β1 integrin inhibits branching morphogenesis. (**A**) Schematics of qPCR primers for testing sgRNA efficiencies. (**B**) Plot of normalized expression levels of wild type *Itgb1* mRNAs by qPCR. The ratio of the testing mRNA to the reference *Rps29* mRNA was normalized to the average value of the sg-Control group. Sample number (left to right): n = 13, 13, 13, 13, 20, 17. (**C**) Confocal fluorescence images of explant cultures transduced with lentiviruses expressing indicated sgRNAs and immunostained with anti-β1 integrin. Explants were fixed at 2 days post transduction. Cyan boxes mark the single-channel magnified images. Asterisks mark cells that are negative for β1 integrin staining but expressing high (yellow asterisks) or low (cyan asterisks) levels of the sgRNA reporter. The NLS-mScarlet channel was subjected to a gamma of 0.5 to facilitate simultaneous display of both bright and dim nuclei. (**D**) Schematic of the quantification method (left) and plots (right) of the β1 integrin intensity across edges of adjacent transduced cells. 5 edges per explant were quantified. Sample number (edges; left to right): n = 15, 15, 25, 25 (day 1); 25, 25, 20, 25 (day 2); 25, 25, 25, 45 (day 4); 25, 25, 25, 25 (day 7). (**E**) Phase contrast images of explant cultures transduced with lentiviruses expressing indicated sgRNAs at 4 different time points (with blasticidin selection). (**F**) Plot of bud number per explant over time. Sample number (left to right): n =19, 19, 19, 20 (day 1); 4, 5, 5, 5 (day 2); 10, 9, 10, 10 (day 3); 15, 15, 15, 15 (day 4); 5, 5, 5, 5 (day 7). Note that only the central image plane of day 7 samples was used for bud counting, whereas all image planes were used for other time points. Error bars in all plots, standard deviation. Statistics, Tukey test (>2 groups) or Student’s t- test (2 groups). *, p<0.05. **, p<0.01. ***, p<0.001. n.s., not significant. Comparisons were all made to the sg-Control group. Scale bars, 20 µm (C), 100 µm (E).

To determine whether and when sgRNA expression could lower protein expression, we performed immunostaining at 1, 2, 4, 7 days after transduction. We found that expressing sg2- or sg3-Itgb1 reduced β1 integrin expression to background levels by day 2 (**Fig. 2C-D**), while a noticeable reduction was observed as early as 1 day post transduction (**Fig. 2D, Appendix Fig. 3A**). On the other hand, expressing sg1-Itgb1 only marginally lowered β1 integrin expression on day 1 (**Fig. 2D, Appendix Fig. 3A**), but the effect reached a similar level as the other two sgRNAs by day 4 (**Appendix Fig. 3A**). Thus, different guide RNAs could have different kinetics when mediating protein reduction. Interestingly, the level of protein reduction was insensitive to the level of sgRNA expression, because cells with low or high sgRNA reporter expression reduced targeted proteins to similar levels (**Fig. 2C**, yellow and cyan asterisks).

We next analyzed explant growth and morphogenesis after gene disruption. All 3 *Itgb1* sgRNAs strongly inhibited the increase in numbers of buds by day 3 (**Fig. 2E-F, Appendix Fig 3B**). The effect of sg1-Itgb1 was less severe than with the other two sgRNAs, consistent with the slower kinetics of sg1-Itgb1 in reducing β1 integrin expression.

To evaluate whether this approach works for salivary epithelial explants starting from a later stage, we performed lentiviral transduction using epithelial bud clusters from E16 Cas9 embryos. We found that expressing any of the 3 *Itgb1* sgRNAs was able to reduce protein expression (**Fig. 3A**) and inhibit budding morphogenesis (**Fig. 3B**). Thus, we conclude that lentivirus-mediated sgRNA delivery can robustly disrupt gene functions in salivary epithelial explants from different embryonic stages.

**Figure 3.**
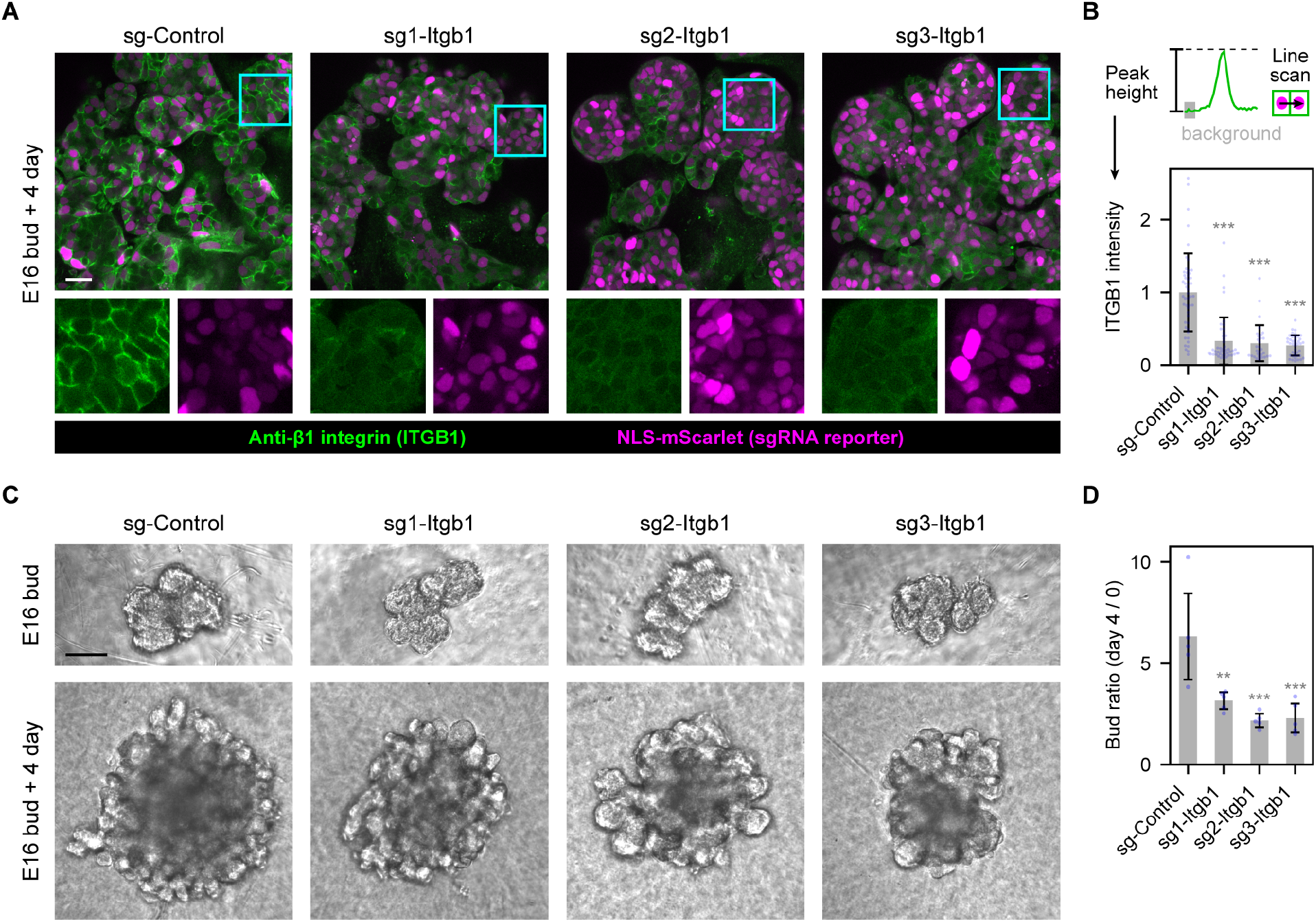
Efficient knockout of β1 integrin in E16 salivary epithelial explants. (**A**)) Confocal fluorescence images of E16 explant cultures transduced with lentiviruses expressing indicated sgRNAs and immunostained with anti-β1 integrin. Explants were fixed at 4 days post transduction. Cyan boxes mark the single-channel magnified images. The NLS-mScarlet channel was subjected to a gamma of 0.5 to facilitate simultaneous display of both bright and dim nuclei. (**B**) Schematic of the quantification method (upper) and plots (lower) of the β1 integrin intensity across edges of adjacent transduced cells. 5 edges per explant were quantified. Sample number (edges; left to right): n = 50, 45, 30, 40. (**C**) Phase contrast images of explant cultures transduced with lentiviruses expressing indicated sgRNAs (with blasticidin selection; 4 days post transduction). (**D**) Plot of bud ratio per explant. Sample number: n = 5 for each group. Note that only the central image plane was used for bud counting. Error bars in all plots, standard deviation. Statistics, Tukey test. **, p<0.01. ***, p<0.001. Comparisons were all made to the sg-Control group. Scale bars, 20 µm (A), 100 µm (C).

### Efficient perturbation of two other genes in salivary epithelial explants

We next tested 3 guide RNAs each for two other genes, *Itga9* and *Serpinh1* (**Appendix Table 1, Appendix Figs. 4A, 5A**). *Itga9* is the most highly expressed α integrin gene in the E13 salivary epithelium (Wang et al. 2021), whereas *Serpinh1* encodes HSP47, a collagen-specific chaperone that is important for collagen biogenesis (Ito and Nagata 2017). The exceptionally high GC content of the *Itga9* sgRNA-containing exon (**Appendix Fig. 4B**) prevented us from qPCR analysis of *Itga9*, whereas qPCR analysis of *Serpinh1* revealed significant reduction of wild type mRNAs when sgRNAs were expressed (**Fig. 5A-B, Appendix Fig. 5B**). For both *Itga9* and *Serpinh1*, we showed that all sgRNAs were able to reduce target protein expression by immunostaining (**Figs. 4A-B, 5C-D**). Therefore, lentivirus-mediated sgRNA delivery in Cas9 transgenic epithelial explants can efficiently reduce protein expression for all three tested genes.

**Figure 4.**
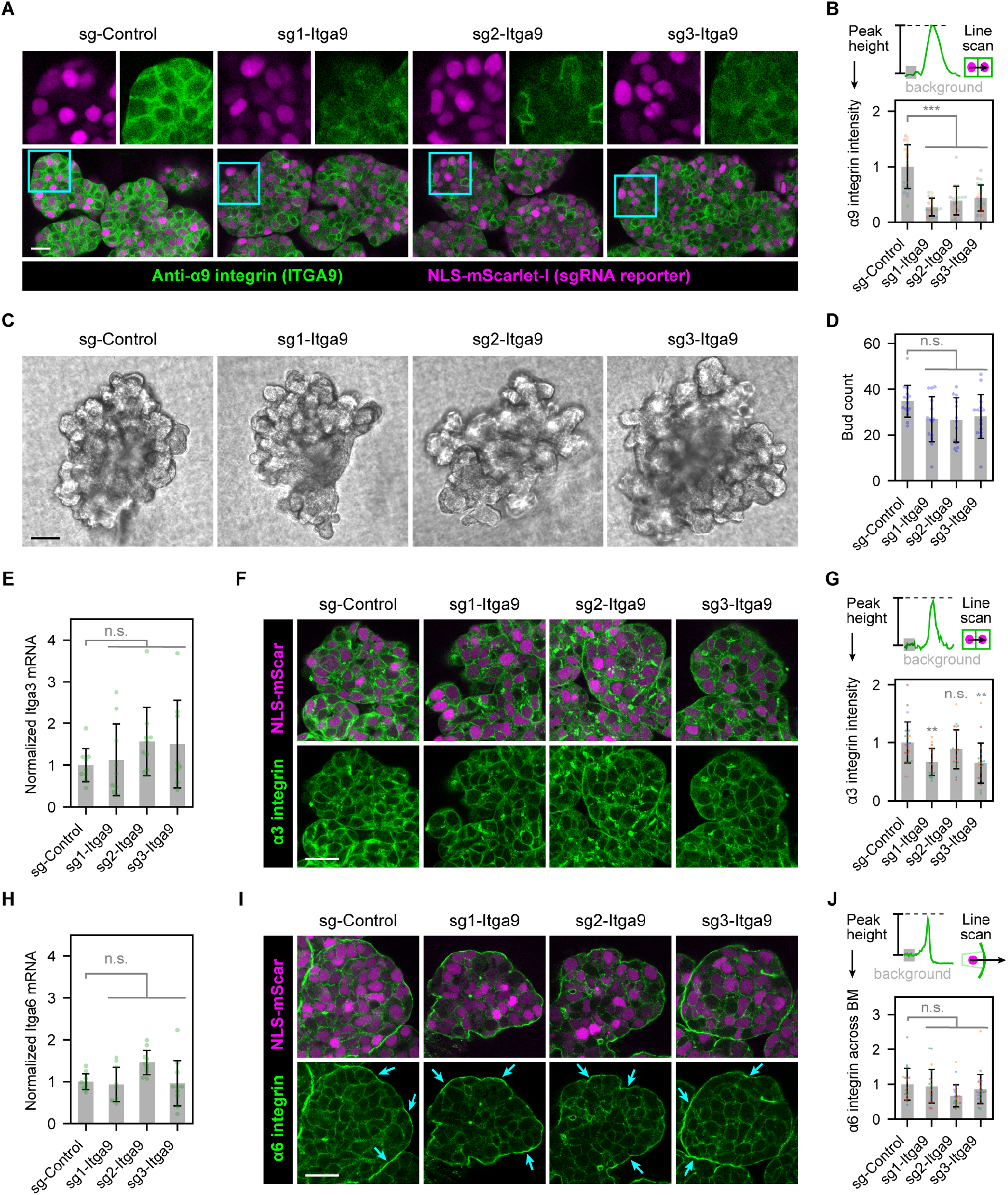
Efficient and specific gene perturbation of *Itga9*. (**A**) Confocal fluorescence images of E13 explant cultures transduced with lentiviruses expressing indicated sgRNAs and immunostained with anti-α9 integrin. Cyan boxes mark the single-channel magnified images shown above. Explants were fixed at 3 days post transduction. The NLS-mScarlet channel was subjected to a gamma of 0.5 to facilitate simultaneous display of both bright and dim nuclei. (**B**) Schematic of the quantification method (upper) and plot (lower) of the α9-integrin intensity across edges of adjacent transduced cells. Sample number: n = 20 edges from 4 different explants for each group. (**C**) Phase contrast images of explant cultures transduced with lentiviruses expressing indicated sgRNAs (with blasticidin selection; 4 days post transduction). (**D**) Plot of bud number per explant. Sample number (left to right): n = 15, 14, 14, 15. (**E, H**) Plot of normalized expression levels of *Itga3* (E) or *Itga6* (H) mRNAs by qPCR. The ratio of the testing mRNA to the reference *Rps29* mRNA was normalized to the average value of the sg-Control group. Sample number (left to right): n = 9, 9, 10, 8 (*Itga3*); 9, 9, 10, 10 (*Itga6*). (**F, I**) Confocal fluorescence images of explant cultures transduced with lentiviruses expressing indicated sgRNAs and immunostained with anti-α3 integrin (F) or anti-α6 integrin (I). Explants were fixed at 4 days post transduction. Cyan arrows point to enriched α6 integrin expression along the bud periphery. The NLS-mScarlet channel was subjected to a gamma of 0.5 to facilitate simultaneous display of both bright and dim nuclei. (**G, J**) Schematics of the quantification method (upper) and plots (lower) of the α3 or α6 integrin intensity. The α3 integrin intensity was quantified across edges of adjacent transduced cells, whereas the α6 integrin intensity was quantified across edges of a transduced cell and the basement membrane. 5 edges per explant were quantified. Sample number (edges; left to right): n = 25, 20, 20, 20 (α3 integrin); 20, 20, 25, 25 (α6 integrin). Error bars in all plots, standard deviation. Statistics, Tukey test. **, p<0.01. ***, p<0.001. n.s., not significant. Scale bars, 20 µm (A, F, I), 100 µm (C).

**Figure 5.**
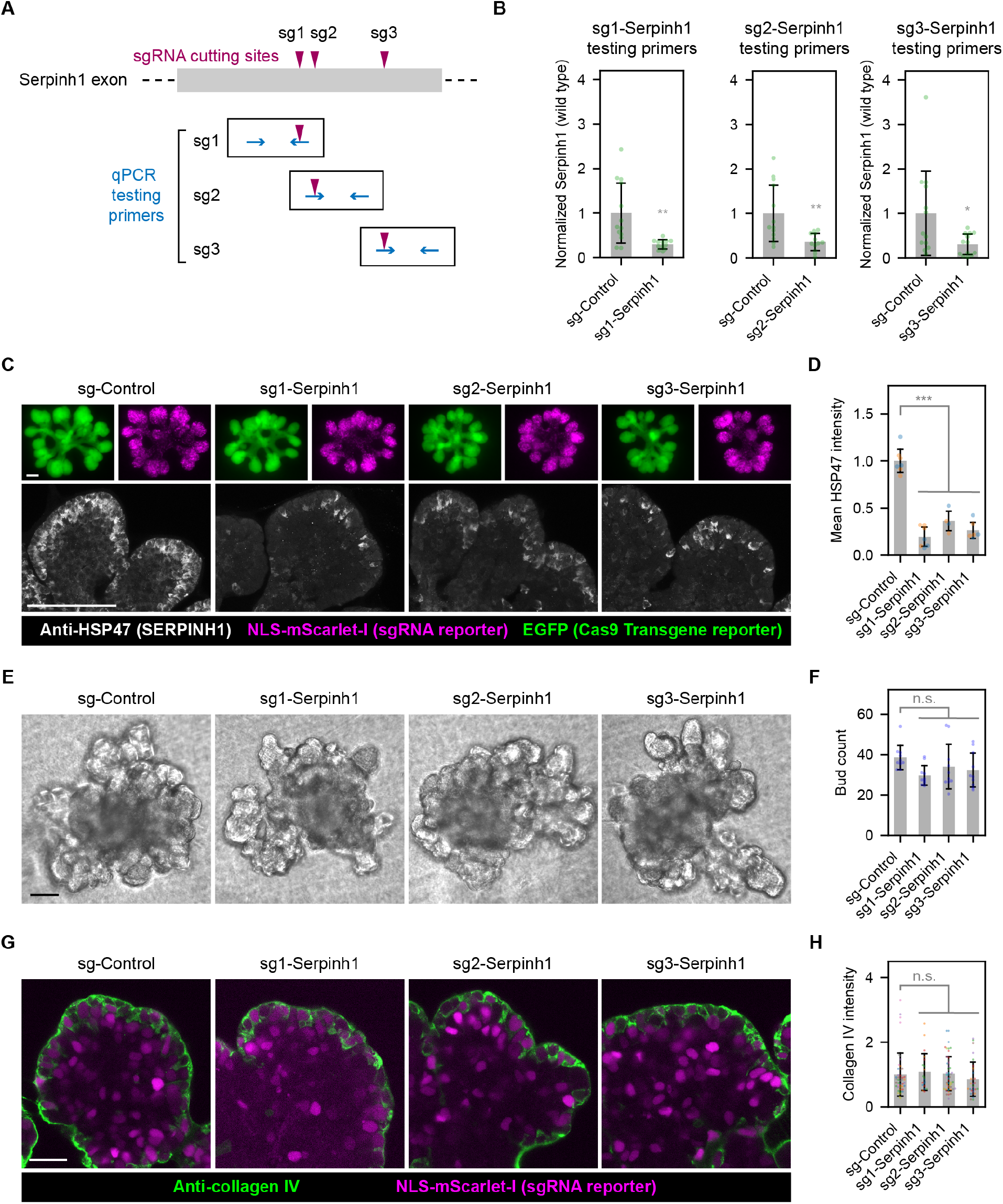
Efficient gene perturbation of *Serpinh1*. (**A**) Schematics of qPCR primers for testing sgRNA efficiencies. (**B**) Plot of normalized expression levels of wild type *Serpinh1* mRNAs by qPCR. The ratio of the testing mRNA to the reference *Rps29* mRNA was normalized to the average value of the sg-Control group. Sample number (left to right): n = 13, 12, 13, 12, 13, 13. (**C**) Widefield (upper) or maximum intensity projection of 10 µm z-range of confocal (lower) fluorescence images of single-bud explant cultures transduced with lentiviruses expressing indicated sgRNAs and immunostained with anti-HSP47 (the *Serpinh1* gene encodes the HSP47 protein). EGFP is co-expressed with the Cas9 transgene. Explants were fixed at 2.5 days post transduction. (**D**) Plot of mean HSP47 intensity per imaging field. Sample number: n = 7, 7, 4, 6 non-overlapping imaging fields from 2 explants for the 4 groups. (**E**) Phase contrast images of explant cultures transduced with lentiviruses expressing indicated sgRNAs (with blasticidin selection; 4 days post transduction). (**F**) Plot of bud number per explant. Sample number: n = 10 for each group. (**G**) Confocal fluorescence images of explant cultures transduced with lentiviruses expressing indicated sgRNAs and immunostained with anti-collagen IV. Explants were fixed at 4 days post transduction. The NLS-mScarlet channel was subjected to a gamma of 0.5 to facilitate simultaneous display of both bright and dim nuclei. (**H**) Plot of the collagen IV intensity across edges between a transduced cell and the basement membrane. Sample number (left to right): n = 45, 20, 50, 30 edges from 3 explants for each group (5 edges per field of view). Error bars in all plots, standard deviation. Statistics, Tukey test (>2 groups) or Student’s t-test (2 groups). *, p<0.05. **, p<0.01. ***, p<0.001. n.s., not significant. Comparisons were all made to the sg-Control group. Scale bars, 100 µm (C, E), 20 µm (G).

Unlike *Itgb1*, no gross phenotypes were observed when *Itga9* or *Serpinh1* were disrupted (**Figs. 4C-D, 5E-F**). We examined the expression of α3 and α6 integrins by qPCR and immunostaining, where we found no or little changes of either upon *Itga9* disruption (**Fig. 4E-J, Appendix Fig. 4C**), suggesting the lack of phenotype when targeting *Itga9* was due to the continued presence of other α integrins that could dimerize with β1 integrin. On the other hand, the lack of phenotype when targeting *Serpinh1* was likely because sufficient collagen IV was provided by the Matrigel supplement. Consistent with this, immunostaining of collagen IV, the major type of collagen in the basement membrane, revealed similar collagen IV expression when Serpinh1 was targeted (**Fig. 5G-H**).

## DISCUSSION

In this study, we describe successful implementation of CRISPR-Cas9 mediated gene knockout in *ex vivo* salivary epithelial explant cultures. We achieved robust gene disruption by transient transduction of high-titer sgRNA-expressing lentiviruses into Cas9-expressing salivary epithelial buds isolated from Cas9 transgenic mice. A major advantage of gene ablation in *ex vivo* cultures is to bypass the need to generate knockout mice to analyze gene functions. To facilitate the application of this method, we established simplified procedures for isolating and culturing salivary epithelial buds that are both less labor-intensive and higher throughput than conventional procedures. We demonstrated successful protein reduction for 3 targeted genes and inhibited branching morphogenesis when targeting the β1 integrin gene. This approach will be broadly useful to analyze the functions of other specific genes using *ex vivo* cultures of various tissues.

### The efficiency of CRISPR-Cas9 mediated gene disruption

While siRNA targets the mRNA, CRISPR-Cas9 targets the genomic DNA. The superior efficacy of CRIPR-Cas9 mediated gene disruption is partly because there are usually only 2 copies of genes, whereas the mRNA molecules of the same gene can number tens to thousands. Moreover, DNA disruption is heritable and thus only needs to occur once, whereas siRNAs need to constantly fight against newly transcribed mRNAs, and they become diluted as cells divide.

The efficiency of CRISPR-Cas9 can be optimized by sgRNA design. Mechanistically, gene disruption by CRISPR-Cas9 mainly comes from frameshift mutations introduced during the error- prone NHEJ repair of DNA breaks. Importantly, the error patterns of NHEJ are strongly influenced by the sequence context. Several studies have developed algorithms that predict how likely an sgRNA will cause out-of-frame mutations (Bae et al. 2014; Doench et al. 2016; Chen et al. 2019). These predictive models have been incorporated as a few efficiency and outcome scores into the intuitive web-based sgRNA design tool CRISPOR to guide the choice of sgRNAs (Concordet and Haeussler 2018).

The lentivirus-based sgRNA delivery ensures persistent sgRNA expression, which further increases the probability of gene disruption. In rare cases, the double-stranded DNA breaks might be repaired perfectly without generating mutations, but these would be cut again until some mutations are generated to prevent further cuts by the CRISPR-Cas9 enzyme. It is worth noting that the U6 promoter we use to drive sgRNA expression imposes two additional considerations on the sgRNA design. First, if the sgRNA does not begin with the nucleotide G, an extra G should be added to the beginning due to transcriptional start site preference of the U6 promoter (Goomer and Kunkel 1992; Ma et al. 2014). Second, certain sequence motifs at the 3’ end of sgRNA can severely lower gene knockout efficiencies (Graf et al. 2019). The sgRNAs containing these motifs are highlighted as “inefficient” when using CRISPOR for sgRNA design, and these sgRNAs should be avoided for gene disruption.

### The specificity of CRISPR-Cas9 mediated gene disruption

Both siRNA and CRISPR-Cas9 have off-target effects, but off-targets of CRISPR-Cas9 can be predicted much more reliably and thus be more effectively mitigated with strategic sgRNA design (Hsu et al. 2013; Bae et al. 2014; Cradick et al. 2014; Doench et al. 2016; Chen et al. 2019). Cutting by CRISPR-Cas9 requires the protospacer adjacent motif (PAM) and is most sensitive to mismatches within 11 bp of the PAM (Jinek et al. 2012; Cong et al. 2013), but it can tolerate certain insertion/deletion mutations in the sgRNA that causes DNA or RNA bulges (Lin et al. 2014). The PAM for the *Streptococcus pyogenes* Cas9 (SpCas9) we use is NGG, which is relatively abundant in the coding sequence of genes. As a result, there are often tens to hundreds of sgRNAs with high-specificity scores from which to choose for any target gene. When selecting sgRNAs for gene knockout studies, it is good practice to prioritize high specificity, and to use several sgRNAs for the same target gene.

### Integrin-mediated cell-matrix adhesion in branching morphogenesis

The importance of β1 integrin in salivary epithelial branching morphogenesis has been established via antibody blocking experiments (Wei et al. 2007). β1 integrin has also been shown to mediate the assembly of human salivary stem/progenitor microstructures by siRNA-mediated knock down (Wu et al. 2019). Our results in this study provide genetic perturbation evidence that strengthens the finding that β1 integrin is required for branching morphogenesis.

β1 integrin can dimerize with multiple types of α integrins to mediate cell-matrix adhesion. We find that disrupting α9 integrin alone does not affect branching morphogenesis, suggesting that either other types of α integrins are more important, or multiple α integrins can act redundantly to provide the necessary cell-matrix interactions for branching morphogenesis.

### Applications of CRISPR-Cas9 gene knockout in *ex vivo* cultures

Lentivirus-based sgRNA delivery for CRISPR-Cas9 gene knockout in *ex vivo* cultures can be easily adapted for analyzing gene functions at the tissue or cell level by adjusting lentivirus titers. For tissue-level phenotypes, such as explant growth and branching morphogenesis, it will be better to use the highest possible titer to transduce most cells in the explant. The penetrance of perturbation phenotypes may be further enhanced by antibiotic selection of transduced cells. For cell-level phenotypes, however, it will be better to lower the virus titer to generate mosaic knockout, so that only a small fraction of cells is perturbed to maintain the native tissue environment. For example, changes of intracellular organization or transcriptional activities during cell differentiation can be examined by immunostaining or single-molecule RNA fluorescence in situ hybridization (smFISH) in mosaic knockout explants (Raj et al. 2008; Wang 2019).

Our work focuses on epithelial cells, but lentivirus can also transduce many other cell types. One possible future application is to test whether it can mediate efficient gene knockout in mesenchymal cells, which will enable studies of specific gene functions in epithelial-mesenchymal interactions.

## AUTHOR CONTRIBUTIONS

R.S. contributed to conception, data acquisition, analysis, drafted and critically revised the manuscript. M.M.M. contributed to data acquisition, and critically revised the manuscript. K.M. contributed to data acquisition, and critically revised the manuscript. S.W. contributed to conception, design, data acquisition, analysis, interpretation, drafted and critically revised the manuscript. All authors gave final approval and agreed to be accountable for all aspects of the work.

## ACKNOWLEDGMENTS

We would like to thank members of the Cell Biology Section for helpful discussions, Kenneth Yamada, and Di Wu (NIDDK) for critical reading of the manuscript, the NIDCR Imaging Core for microscopy support, the NIDCR Combined Technical Research Core for Sanger sequencing, and the NIDCR Veterinary Resource Core for animal support. S.W. is supported in part by an NIDCR K99 Pathway to Independence Award (K99 DE27982). This work is supported by the NIH Intramural Research Program (NIDCR, ZIA DE000525). The authors declare that there is no conflict of interest.

---

**Appendix Fig. 1.**
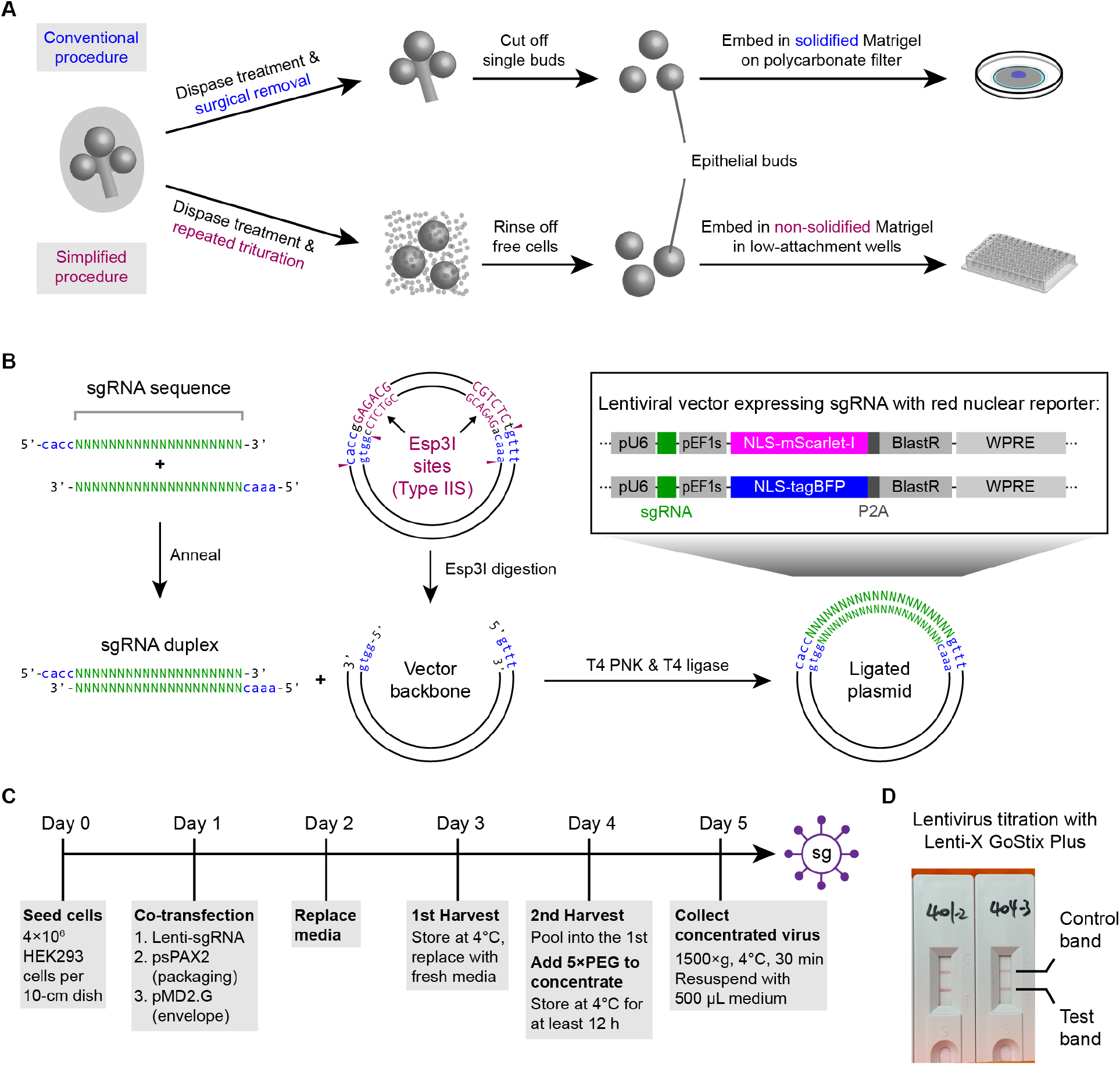
(Related to Fig. 1). (**A**) Schematics showing conventional (upper) and simplified (lower) procedures to isolate and culture single epithelial buds from mouse embryonic salivary glands. (**B**) Schematics showing cloning procedure for the sgRNA-expressing lentiviral vector. (**C**) Schematics showing the timeline of lentivirus production. (**D**) Image of lentivirus titration results using Lenti-X GoStix Plus.

**Appendix Fig. 2.**
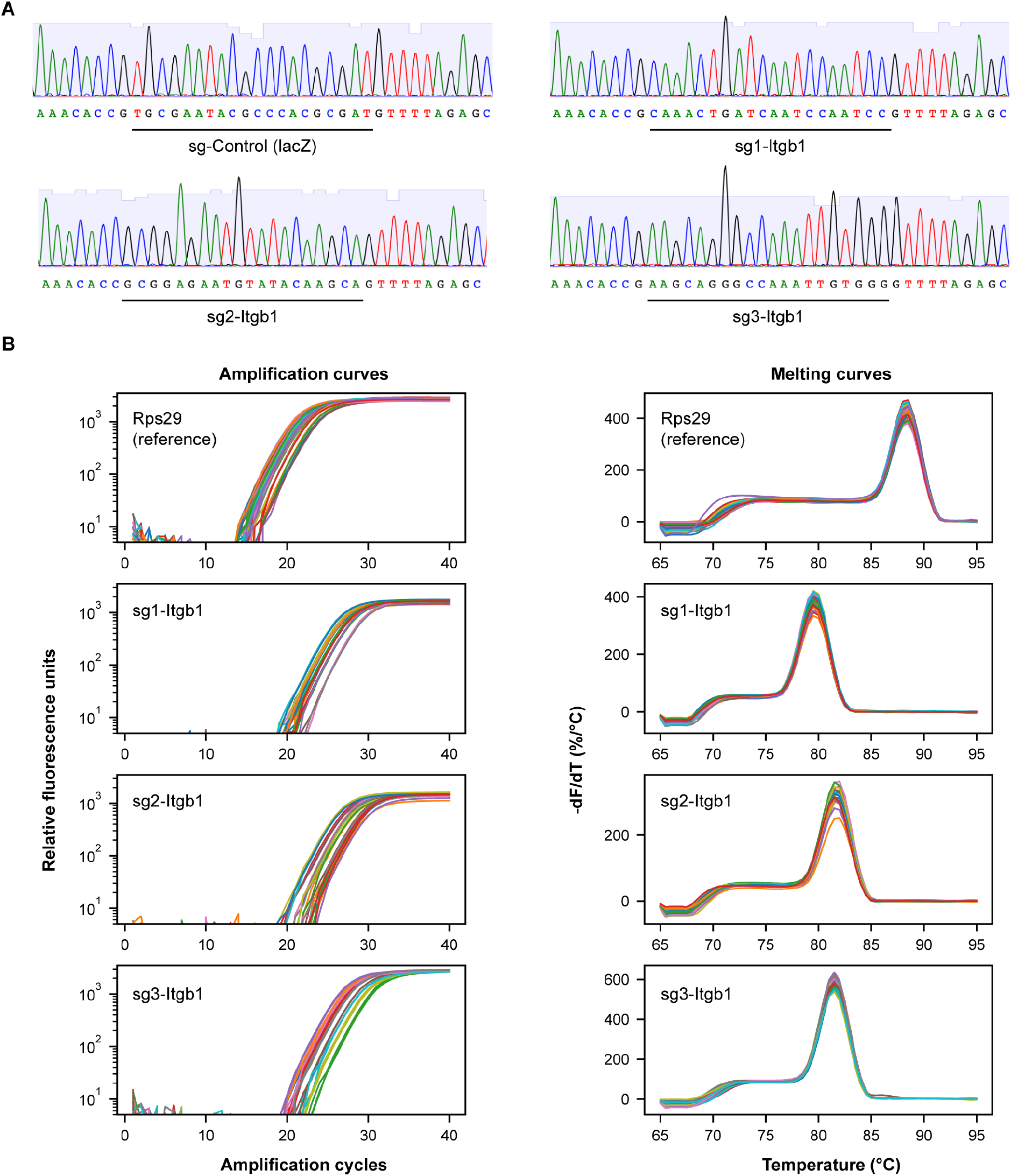
(Related to Fig. 2). (**A**) Sanger Sequencing verification of indicated sgRNA sequences. (**B**) Amplification curves and melting curves of indicated qPCR primers.

**Appendix Fig. 3.**
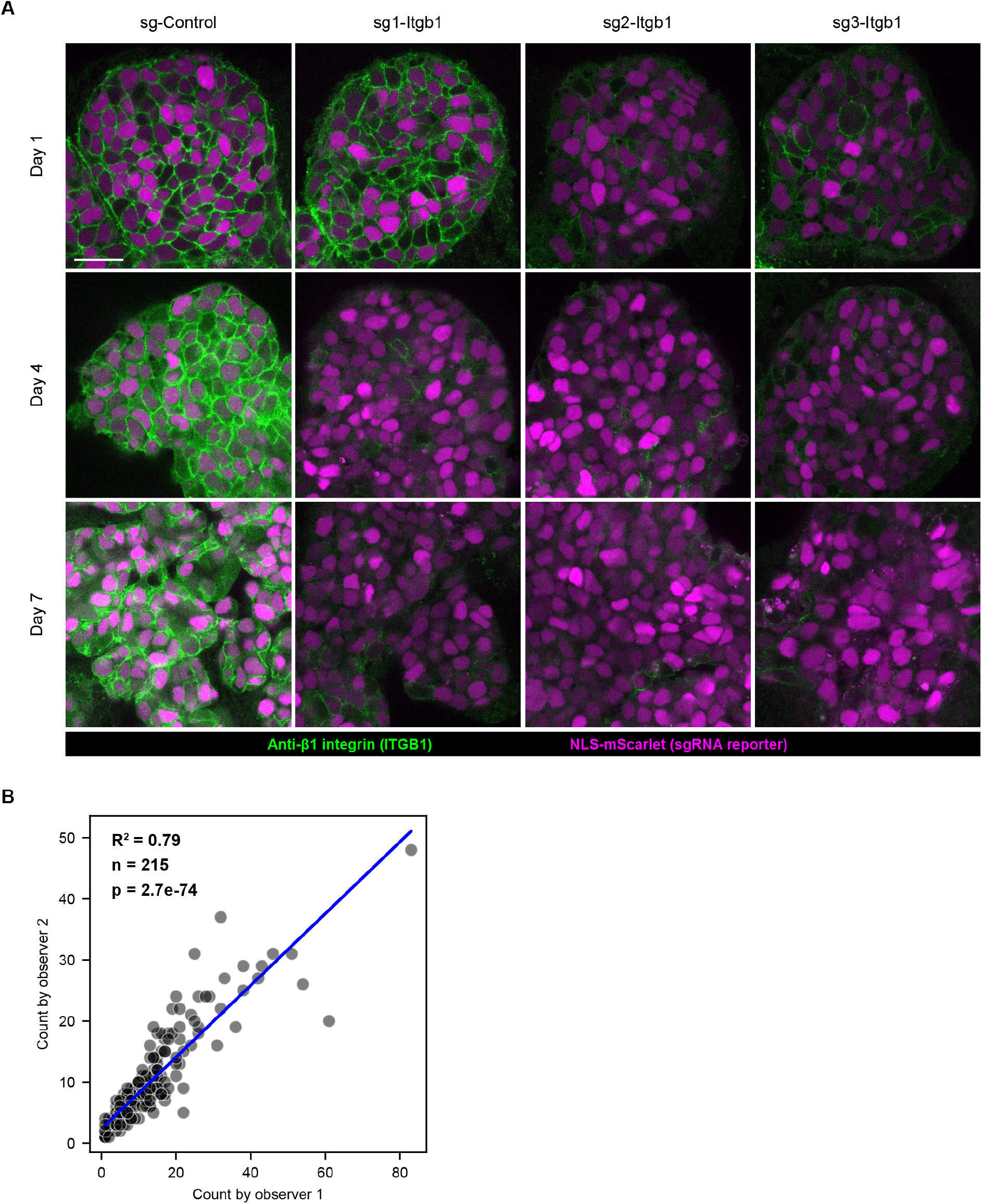
(Related to Fig. 2). (**A**) Confocal fluorescence images of explant cultures transduced with lentiviruses expressing indicated sgRNAs, fixed at indicated days post transduction, and immunostained with anti-β1 integrin. The NLS-mScarlet channel was subjected to a gamma of 0.5 to facilitate simultaneous display of both bright and dim nuclei. Scale bar, 20 µm. (**B**) Scatterplot showing the positive correlation of bud counting by two observers. Blue line represents the linear regression.

**Appendix Fig. 4.**
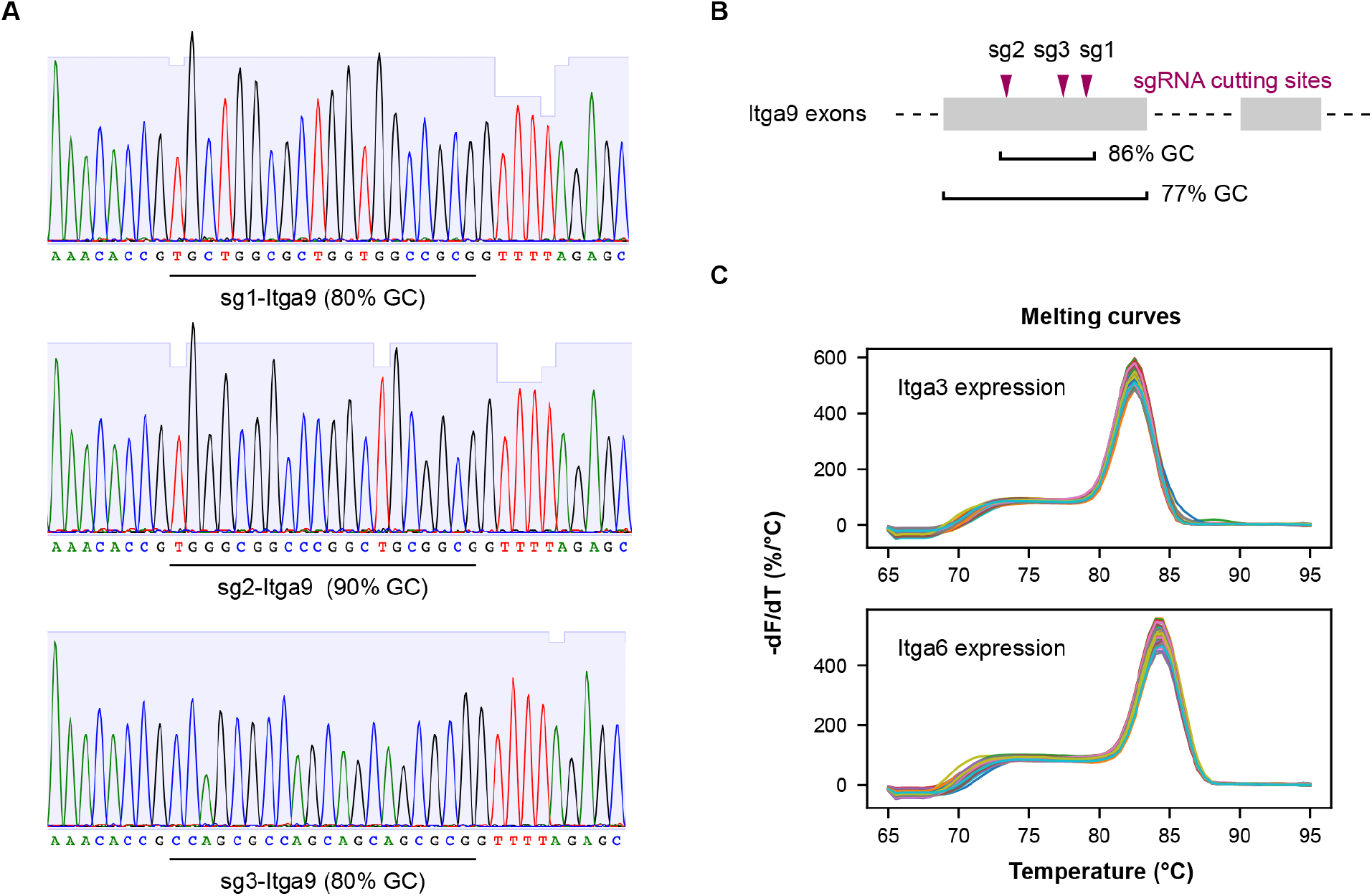
(Related to Fig. 4). (**A**) Sanger sequencing verification of sgRNAs targeting *Itga9*. (**B**) Schematic summarizing the GC content in the first exon of *Itga9*, which contains the targeted sites of all 3 sgRNAs. (**C**) Melting curves of qPCR primers for testing *Itag3* and *Itga6* expression.

**Appendix Fig. 5.**
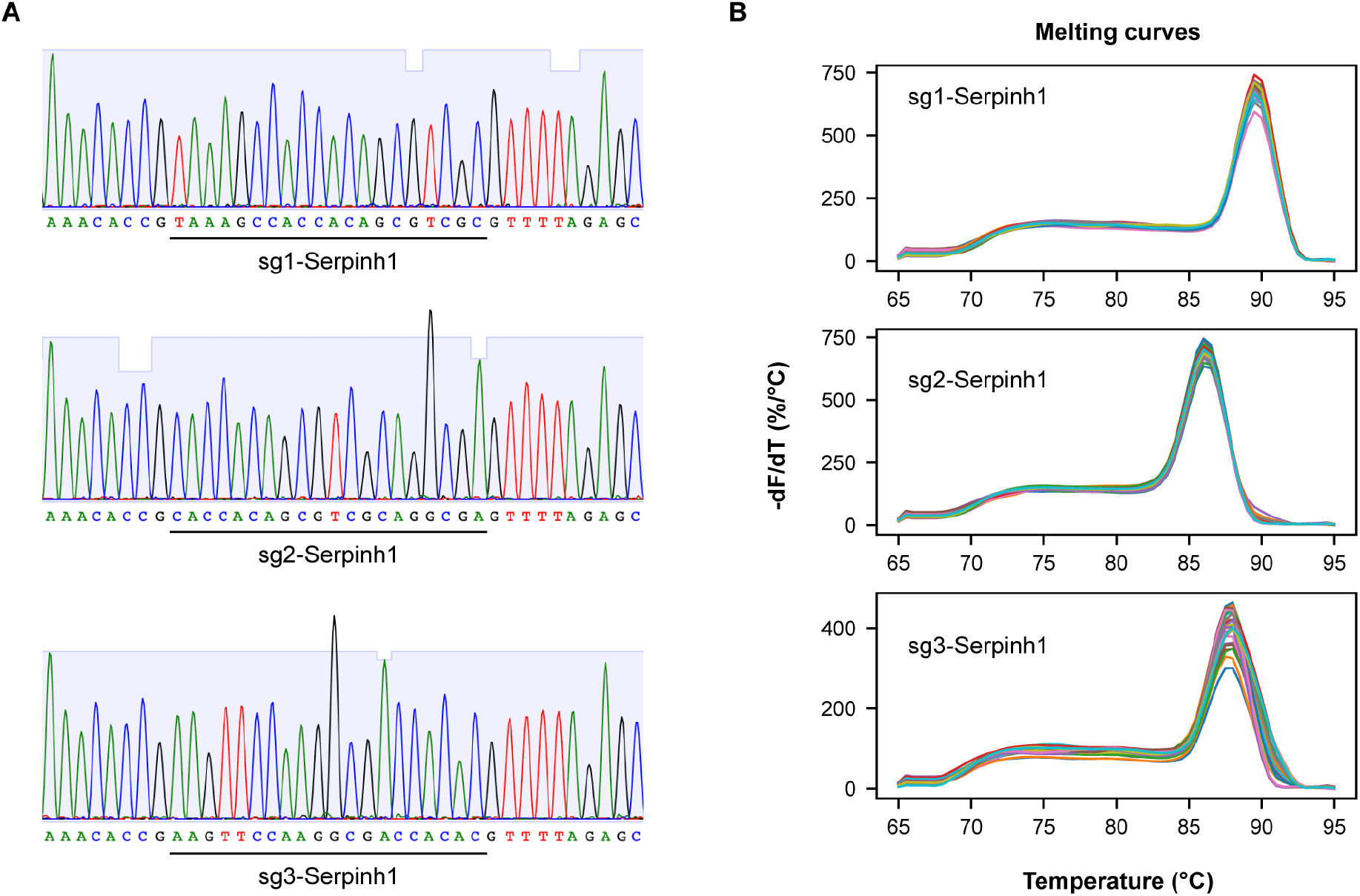
(Related to Fig. 5). (**A**) Sanger sequencing verification of sgRNAs targeting Serpinh1. (**B**) Melting curves of qPCR primers for testing 3 sgRNAs targeting Serpinh1.

**Appendix Table 1.**
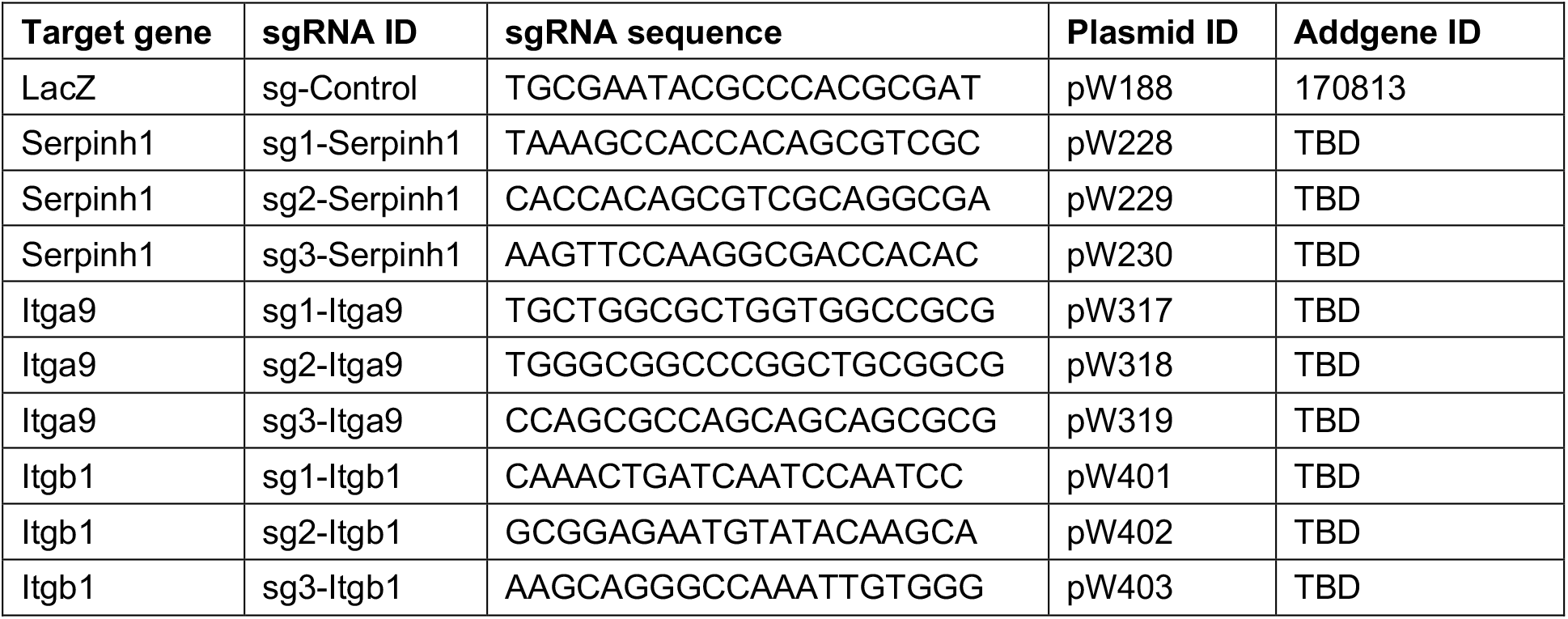
sgRNA sequences.

**Appendix Table 2.**
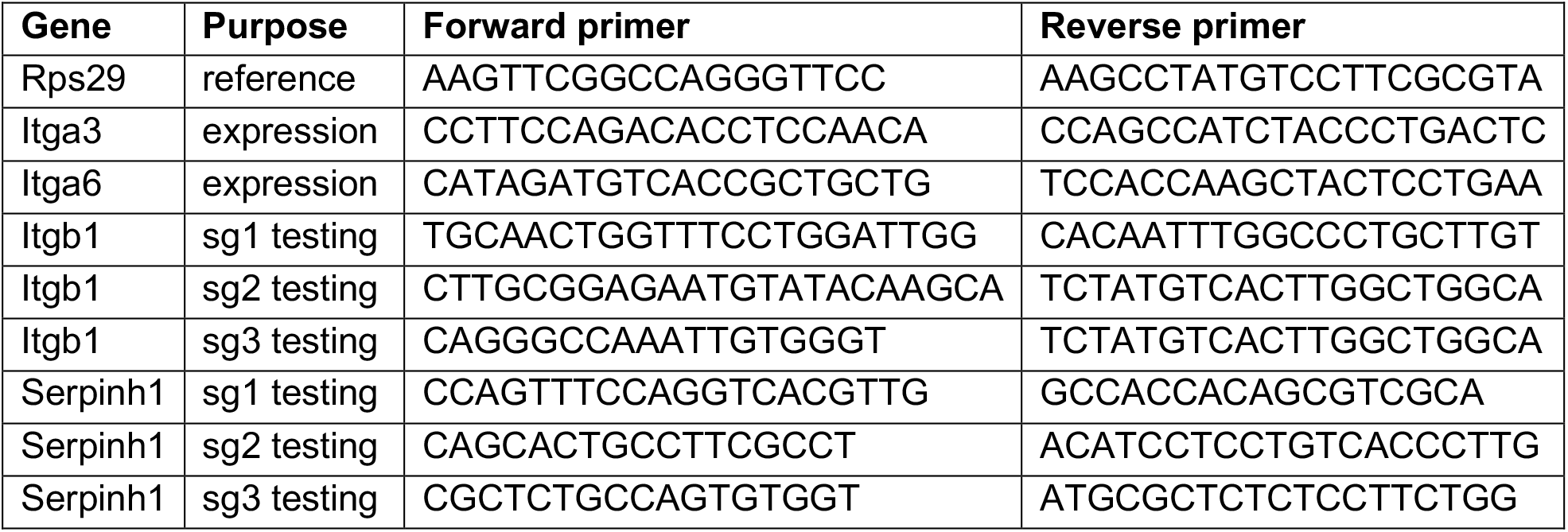
qPCR primers.

## Notes

### Competing Interest Statement

The authors have declared no competing interest.

### Summary of Updates

Figures expanded and revised to add new data supporting the effectiveness of the method

